# Designing Cost-Effective, Open-Source, Multi-Head Bioprinters via Conversion of Hobby-Grade 3D Printers

**DOI:** 10.1101/2022.03.24.483055

**Authors:** David Chimene, Kaivalya A. Deo, Jeremy Thomas, Akhilesh K. Gaharwar

## Abstract

Over the past decade, additive manufacturing has resulted in significant advances towards fabricating anatomic-size, patient-specific scaffolds for tissue models and regenerative medicine. This can be attributed to the development of advanced bioinks capable of precise deposition of cells and biomaterials. The combination of additive manufacturing with advanced bioinks is enabling researchers to fabricate intricate tissue scaffolds that recreate the complex spatial distributions of cells and bioactive cues found in the human body. However, the expansion of this promising technique has been hampered by the high cost of commercially available bioprinters and proprietary software. In contrast, conventional 3D printing has become increasingly popular with home hobbyists and caused an explosion of both low-cost thermoplastic 3D printers and open source software to control the printer. In this work, we bring these benefits into the field of bioprinting by converting widely available and cost-effective 3D printers into fully functional, open source, and customizable multi-head bioprinters. We demonstrate the practicality of this approach by designing bioprinters customized with multiple extruders, automatic bed leveling, and temperature controls for approximately $400. These bioprinters were then used for in vitro and ex vivo bioprinting to demonstrate their utility for tissue engineering.

## 1. INTRODUCTION

Bioprinting represents an important and growing branch of tissue engineering because it allows researchers to deposit cells and biomaterials in precise distributions that mimic native tissue architectures. Although initially bioprinting was restricted to small, flat structures by the lack of suitable bioinks able to combine high print performance with cell viability, its potential nevertheless sparked a rapid growth in popularity over the last decade.^1–2^ This increased attention from tissue engineers and materials scientists led to breakthroughs in high performance bioinks that enabled taller structures to be bioprinted that are reminiscent of true 3D printing. These advanced bioinks are created primarily from cell friendly hydrogels that are efficiently imbued with improved flow properties and more robust mechanical properties through a variety of methods.^3^ Advanced bioinks allow the creation of full scale tissue reconstructions including of bone,^4–5^ cartilage,^6^ vasculature,^7^ and skin.^8^ More bioinks are also under development that maintain a microenvironment containing cues that induce stem cell proliferation & differentiation, and that can be enzymatically remodeled by patient stem cells over time.^4, 9^ This is leading into research on in vivo implantable tissue scaffolds, and promises that bioprinting will soon enable rapid fabrication of implantable human tissue constructs that can remodel into healthy functional tissue at the implantation site. By allowing patient tissue to be regrown, this technology holds the potential to revolutionize medicine through treatments ranging from bone & skin reconstruction to diabetes, arthritis, and cardiovascular disease.^4–8^

The enormous promise of bioprinting has naturally led to rapidly increasing interest from the research community, and advanced bioinks have markedly increased the capabilities of bioprinting.^10^ However, one key impediment to increasing research in this promising field is the high price of entry. Bioprinters on the market today range from $10,000 over $1,000,000.^2, 5, 11^ This represents a significant impediment to most research labs entering into this research field. Additionally, commercial bioprinters run on limited proprietary software, which stifles research innovation by preventing customizations that may be necessary to pursue new research ideas.

Not long ago, 3D printing was also limited to expensive commercial machines and proprietary software, as bioprinting is today. However, once the Stratasys patent on fused deposition modeling expired in 2009, the price of 3D printers dropped dramatically from over 10,000$ to under 1,000$. It also saw the rapid development of community-created open source software that enabled printing enthusiasts to develop their own tools for improving their printers as they saw fit. Today, home 3D printers start at under 200$.^12^ These changes have democratized 3D printing and caused an explosion of popularity, with over a million 3D printers estimated to have been sold in 2020.^13^

Here we aim to combine the low-cost printers and open source tools of traditional 3D printing into the field of bioprinting. This will lower the costs of entry into bioprinting research and encourage innovation, helping to fuel research into this promising technology. These effects occurred in the 3D printing world because of the thousands of dedicated enthusiasts creating, maintaining, and sharing code with each other. Fortunately, this is not necessary to bring these advances to the field of bioprinting. Because of the many similarities between bioprinters and thermoplastic 3D printers, much of the low-cost hardware and open source software of 3D printers can be tweaked slightly to create an open source bioprinter that can be created for about 400$ (**Figure 1**).

**Figure 1.**
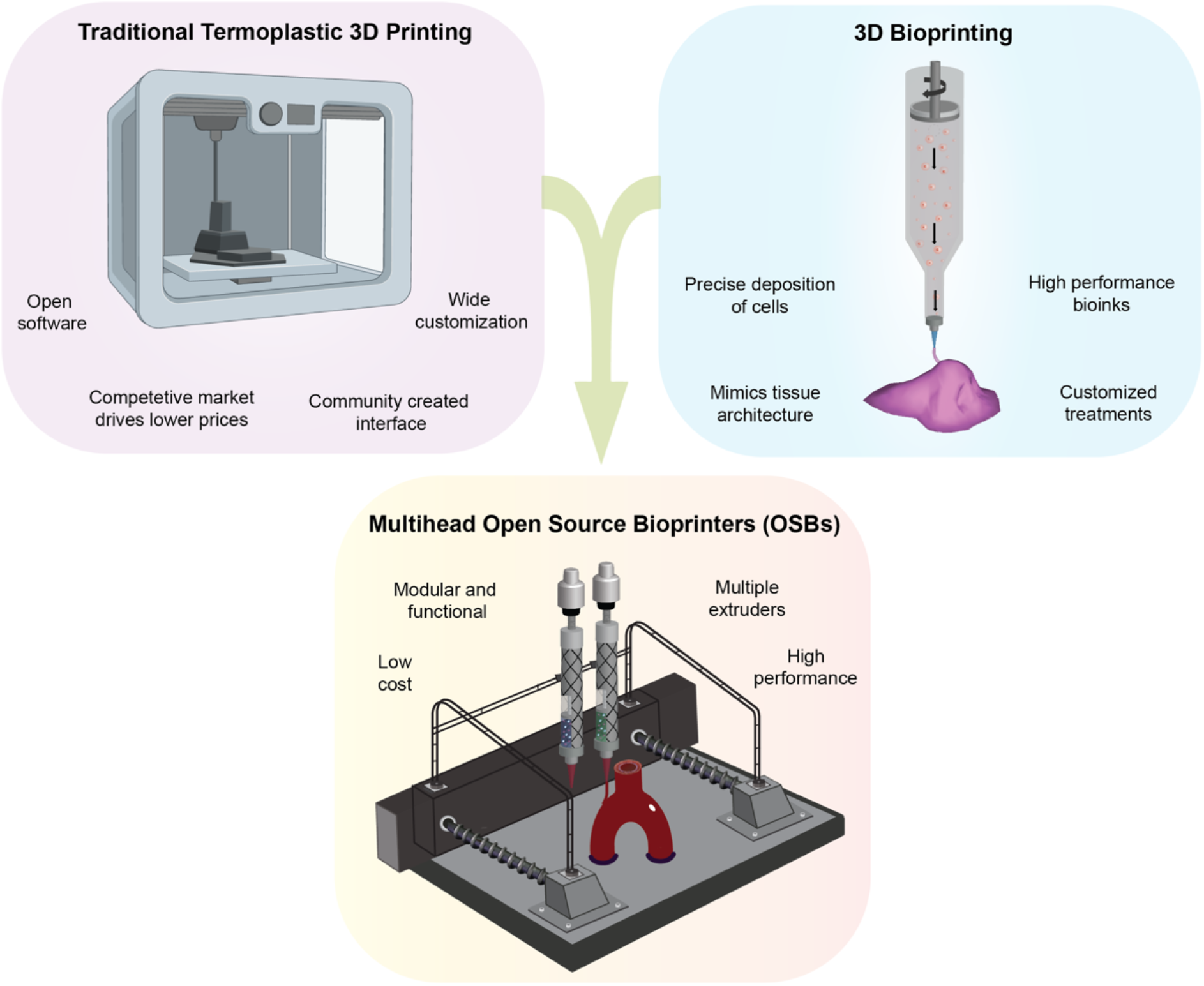
Engineering Multihead Open Source Bioprinters (OSBs). Building functional, open source 3D Bioprinters from low-cost thermoplastic 3D printers enabling accessibility to high performance 3D bioprinting technology. The OSBs are highly customizable with options of automatic bed leveling, multiple extruders and bed/extruder temperature control.

The high cost and restrictive software of existing bioprinters has led other labs to publish papers on bioprinter construction in the last year. Lanaro et al. created an impressive biofabricator that combined electrospinning with pneumatic extrusion.^14^ However, this machine costs over 10,000$ to build in parts alone and requires custom machined parts. Likewise, Yenilmez et al. developed a hybrid inkjet-coaxial pneumatic extrusion bioprinter using custom made syringe pumps and controller on a CNC stage.^11^ These papers demonstrate methods of building a bioprinter from scratch and both require custom parts to be machined, which restricts the number of labs that can reasonably adapt these prototypes to their needs. In contrast to the complexity of building a printer from the ground up, in this paper we demonstrate a much simpler method by converting an existing mass-produced thermoplastic 3D printer into a bioprinter, using the 3D printer itself to fabricate the necessary parts. This conversion can be done with little 3D printer specific expertise and does not require machining parts, allowing biomaterials-focused research groups to easily create their own bioprinters. In addition, this design costs much less than those described above, approximately $400, depending on the chosen customizations.

Finally, this paper shows for the first time how to create an open source, multi-head extruder that uses motor-driven extrusion. The ability to customize and append multiple extruders enables multi-material 3D bioprinting, permitting better recapitulation of tissue microarchitecture. Additionally, this extrusion modality is more suitable than pneumatic extrusion for advanced bioinks with complex rheological properties because it allows volumetric extrusion to occur regardless of rheological properties like thermal gelation, shear thinning, yield stress, and thixotropic effects. These complex rheological properties are commonly seen in newer advanced bioinks, where these flow properties are exploited to improve print performance.^3, 15^

We demonstrate our bioprinters’ practical utility by using them for 3D bioprinting tests incorporating murine preosteoblast cells into an advanced nanoengineered ionic covalent entanglement (NICE) bioink optimized for bone regeneration **(Figure 8a)**.^4, 16–17^ We also used this bioprinter to print another NICE bioink formulation directly into the cartilage of an ex vivo equine meniscus **(Figure 8b)**.

In this paper, we demonstrate how to convert one of the most popular hobbyist 3D printers, the Ender 3 Pro, into a fully functional bioprinter running on open source tools and freeware. This conversion procedure is also applicable to most common thermoplastic 3D printers, allowing researchers to use this method to easily build bioprinters customized to their own research needs. The 3D printer itself is first used to print out hardware components for its own conversion. Then, the hardware is switched over along with some easily sourced components to create the bioprinter hardware. Next, we show how to install open source firmware onto the printer control board, along with key tweaks to make it function as a bioprinter. Finally, we discuss how to use open source 3D printer software to operate the bioprinter. We also cover examples of several bioprinter customization options, including multiple-extruder designs, temperature-controlled extruder barrels, and automatic bed leveling.

## 2. RESULTS AND DISCUSSION

This section contains descriptions of the hardware and firmware conversion processes used to convert a thermoplastic 3D printer into a bioprinter. Because this paper is focused on creating an easily customizable platform, instructions for several options are listed, including automatic bed leveling options, multiple extruders, and temperature controls. Most of these options require only small variations of the same conversion process, so the additional firmware and hardware steps are listed separately without repeating the bulk of the instructions. Only the dual extrusion option diverges enough to warrant a separate set of steps. For each of the options listed, the finished configuration files and 3D models are included in the supplementary data. Interested researchers would simply need to print out the appropriate hardware parts and upload the attached firmware in order to complete the bioprinter conversion.

### 2.1 Initial 3D Printer Choice & Setup

This conversion was based around the Ender 3 Pro, a ubiquitous 3D printer model that retails for 209$ and combines an all-metal frame with a small footprint, making it easy to transport and sterilize. This conversion will also work with many other common thermoplastic printer models with minor changes to the firmware. This conversion can be applied to most common thermoplastic 3D printers, allowing researchers to customize their open source bioprinter to meet their research needs. For example, this procedure can be applied with only minor firmware changes to create bioprinters with larger print beds, larger motors for faster printing, and additional extruders beyond the dual extrusion demonstrated here.

The first step in any bioprinter conversion is to set up the 3D printer and use it to print copies of the appropriate conversion hardware from **Table 1 (Figure 2a-c)**, including a syringe bracket assembly **(Figure 2a)**, a motor mount **(Figure 2b)**, and a motor adapter **(Figure 2c)**. These are combined with a NEMA 11 19:1 geared stepper motor and a MGN12 linear rail with a compatible MGN12H carriage block to convert the bioprinter. A complete list of materials is attached in **Table 2-4** and 3D printer parts **(Supplementary file)**.

**Table 1.**
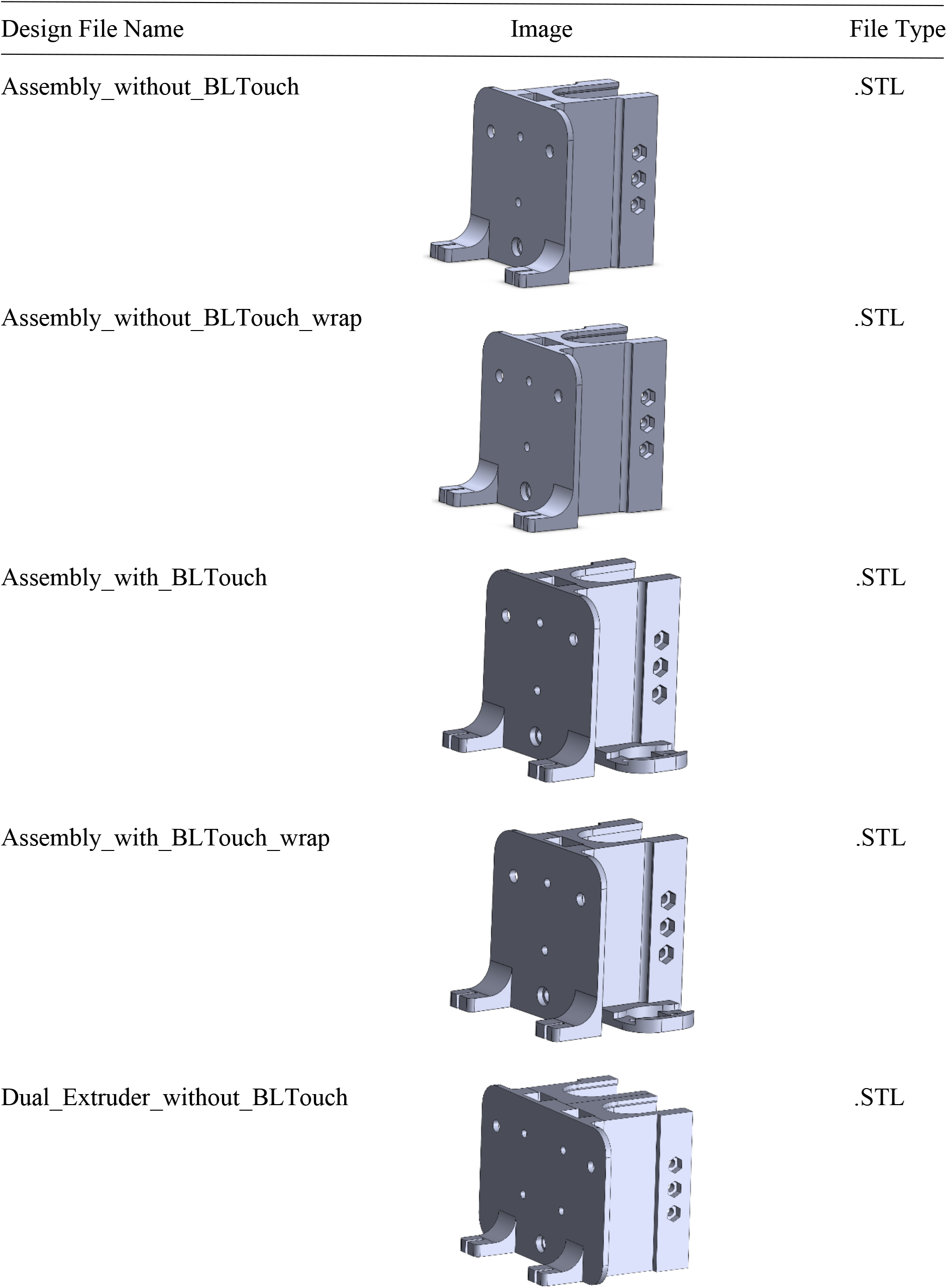
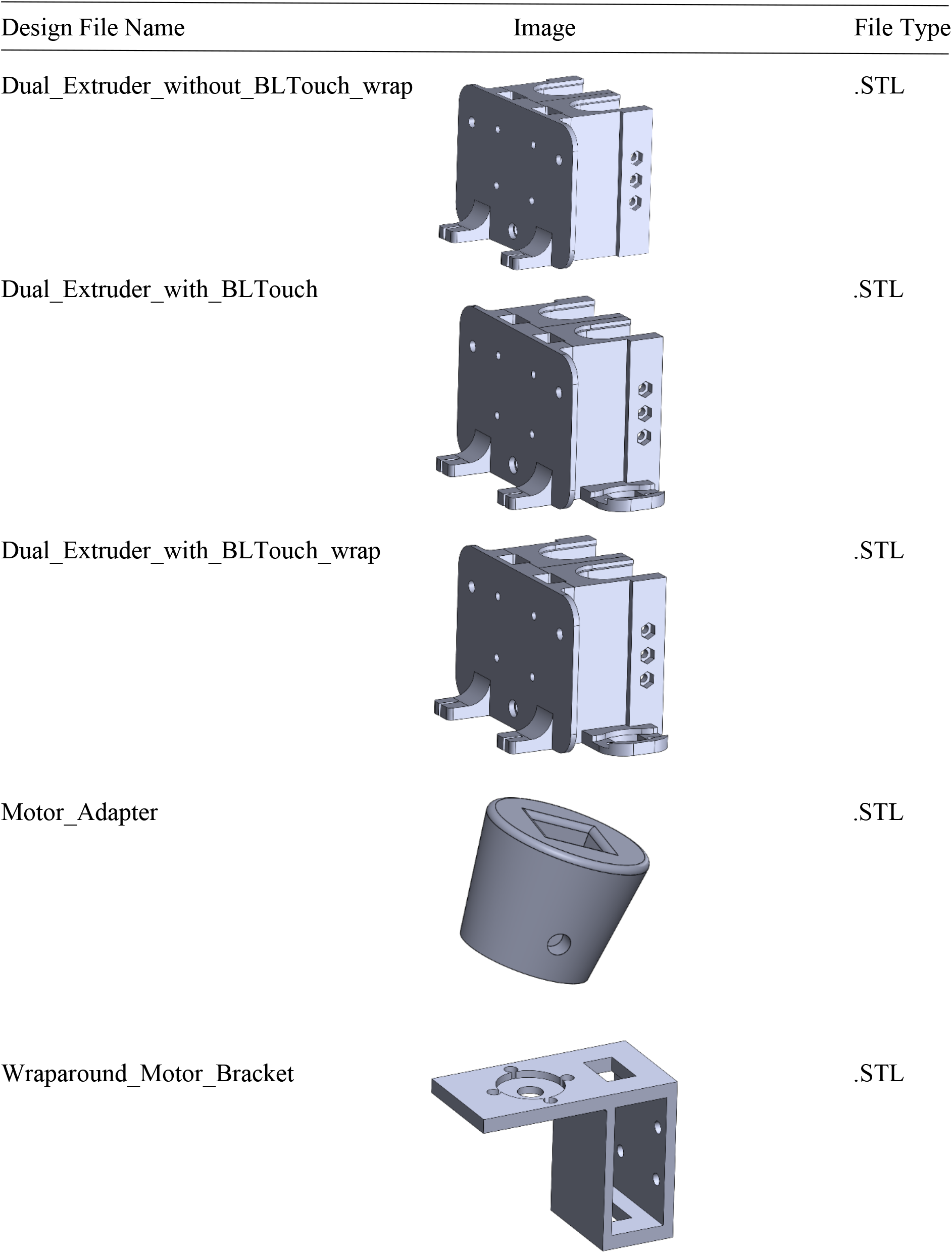
Design Files Summary

| Design File Name | Image | File Type |
| --- | --- | --- |
| Assembly_without_BLTouch |  | .STL |
| Assembly_without_BLTouch_wrap |  | .STL |
| Assembly_with_BLTouch |  | .STL |
| Assembly_with_BLTouch_wrap |  | .STL |
| Dual_Extruder_without_BLTouch |  | .STL |

*(continued on next page)*

**Table 1** (*continued*)
| Design File Name | Image | File Type |
| --- | --- | --- |
| Dual_Extruder_without_BLTouch_wrap |  | .STL |
| Dual_Extruder_with_BLTouch |  | .STL |
| Dual_Extruder_with_BLTouch_wrap |  | .STL |
| Motor_Adapter |  | .STL |
| Wraparound_Motor_Bracket |  | .STL |

**Figure 2.**
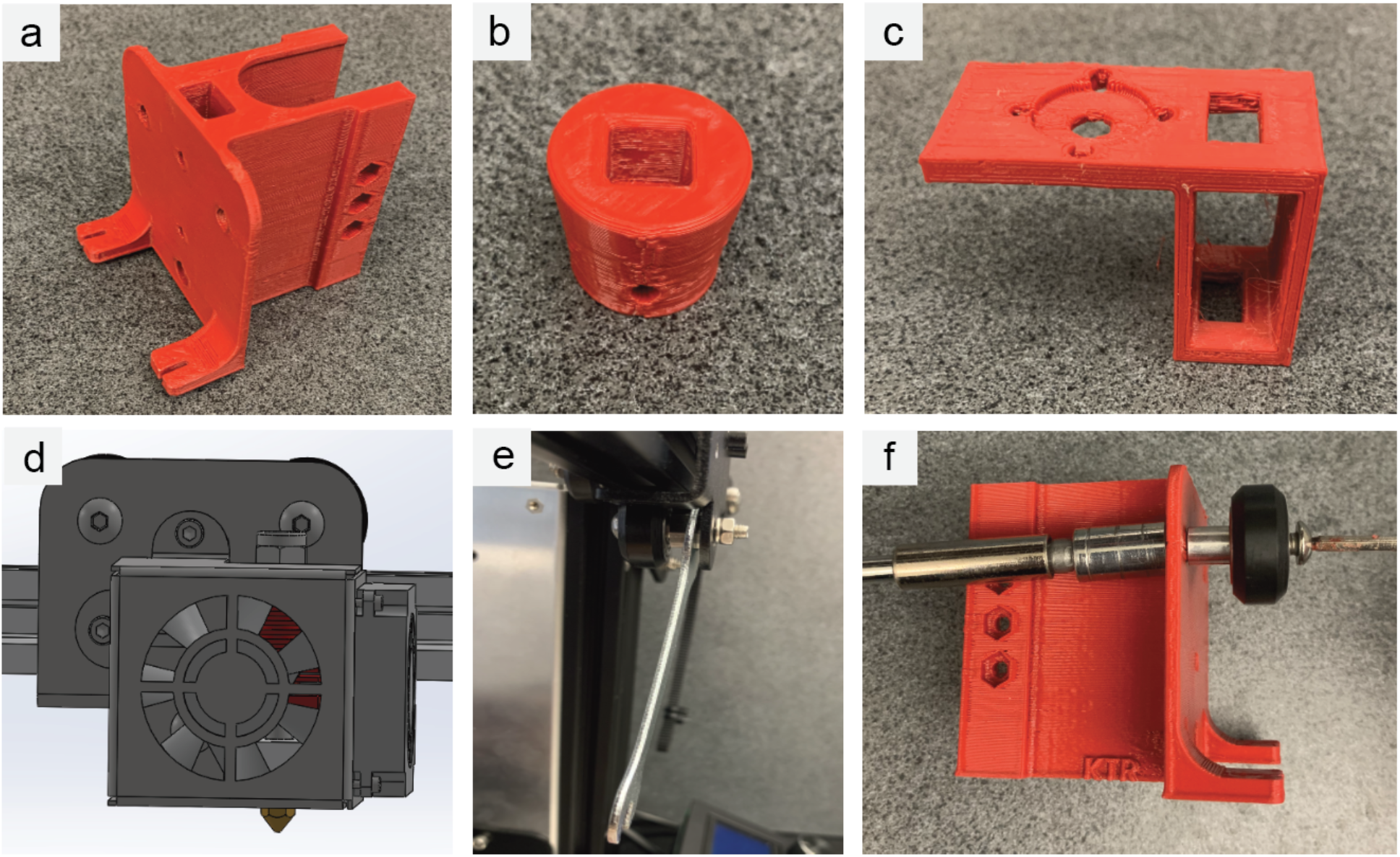
Hardware Part Conversion-I. (a) The syringe bracket assembly interfaces with the x gantry and holds the syringe in place during bioprinting. It also serves as a connection point for the linear rail. (b) The motor adapter is designed to translate torque from the two geared stepper motor to the syringe. (c) The motor mount secures the motor to the linear rail carriage block. (d) The thermoplastic extruder hardware pictured here is removed, including the fan shroud, hot end, and extruder backplate. (e) The extruder backplate can be removed from the x gantry by loosening the eccentric nut on the lower wheel. (f) The wheels are transferred from the extruder backplate onto the syringe bracket assembly.

**Table 2.** Main Printer Supplies.

| Item | Price |
| --- | --- |
| *Ender 3 Pro* | \$209.00 |
| *19:1 Planetary Gearbox NEMA 11 Stepper Motor*<br>*Extruder 3D Printer Motor* | \$40.49 |
| *ReliaBot 200mm MGN12 Linear Rail Guide with*<br>*MGN12H Carriage Block for 3D Printer and CNC*<br>*Machine* | \$26.29 |
| *Makin's USA Professional Ultimate Clay Gun Extruder* | \$36.52 |

**Table 3.** Optional Parts.

| Item | Price |
| --- | --- |
| *EZABLTM Pro Mini– Auto Bed Leveling Kit w/ OEM*<br>*Ender 3 Mount* | \$58.99 |
| *ANTCLABS BLTouch : (With 1M Extension Cable Set)* | \$39.29 |
| *Pin 27 Adapter Board* | \$10.99 |
| *Steel Build Plate 235mm x 235mm* | \$14.99 |
| *USB to Mini-USB data cable* | \$4.79 |
| *Uno Bootloader Flashing Kit(Arduino Uno, USB 2.0*<br>*Cable, M-F & F-F Dupont Wires)* | \$15.99 |

**Table 4.** Recommended Tools.

| Item | Price |
| --- | --- |
| *Calipers* | \$19.99 |
| *M3 Hex Head Nut & Bolts assortment pack* | \$16.99 |
| *Wire Crimpers* | \$13.99 |
| *JST-XH Connector Kit with 2.54mm Female Pin Header*<br>*and 2/3/4/5/6 Pin Housing* | \$9.29 |
| *Size 111 Viton O-Ring* | \$8.59 |
| *Size 007 Viton O-Ring* | \$5.53 |

### 2.2 Hardware Conversion

The first step in hardware conversion is removal of the thermoplastic extrusion hardware. This includes removing the fan shroud and hot end, as seen in **(Figure 2d)**. Next, the extruder backplate is removed from the x-gantry using a wrench to loosen the eccentric nut **(Figure 2e)**. The wheels from the extruder backplate are then transferred to the syringe bracket assembly **(Figure 2f)**.

Then, the geared stepper motor is attached to the motor mount **(Figure 3a)**, and the motor mount is bolted to the carriage block to secure it to the linear rail **(Figure 3b)**. An M3 nut and bolt are inserted into the motor adapter and tightened onto the motor shaft **(Figure 3c)**. The linear rail is then inserted into the syringe bracket assembly **(Figure 3D)**.

**Figure 3.**
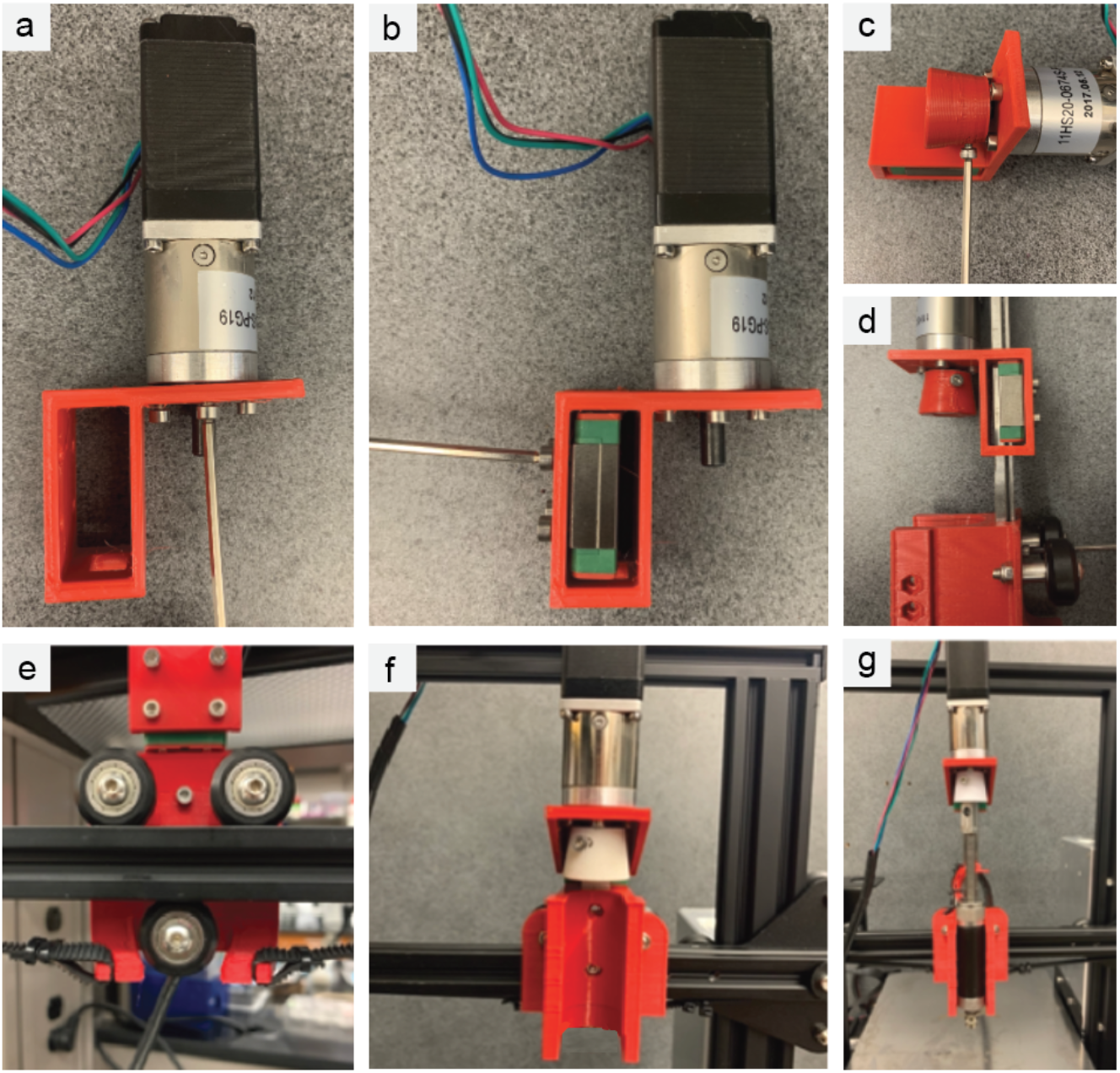
Hardware Part Conversion-II. (a)The geared stepper motor was attached to the motor mount using 4 m^3^ screws. (b) The motor mount was then attached to the linear rail carriage block using 4 m^3^ screws. (c) The motor adapter was then affixed to the motor shaft using 1 m^3^ screw and nut. The linear rail was then inserted into the syringe bracket assembly using 2 m^3^ screws and nuts. (d) The syringe bracket assembly was slid onto the x-gantry and tightened in place using the eccentric nut on the lower wheel, and the x-axis belt was attached to the belt slots. (e) The completed bioprinter assembly, including SBA, linear rail, motor mount, geared stepper motor, and motor adapter. (f) The extruder syringe should fit tightly into the bracket and motor adapter, allowing the printer to control extrusion using the stepper motor.

Next, the syringe bracket assembly is mounted to the x-gantry in place of the original extruder backplate by tightening the eccentric nut on the lower wheel to hold it in place. Then, the x-axis belt is attached in the belt slots on the syringe bracket assembly **(Figure 3e)**. Once this is complete, the syringe bracket assembly together with the linear rail, carriage block, motor mount, motor, and motor adapter all move together as one unit across the x gantry, just as the extruder backplate, hot end assembly, and fan shroud do on the thermoplastic 3D printer **(Figure 3f)**. The syringe barrel can then be inserted into the syringe bracket assembly and secured into the motor adapter, allowing the motor to control extrusion **(Figure 3g)**. Finally, the geared stepper motor is plugged into the extruder motor connection on the main board **(Figure 4a)**, and the voltage is adjusted to the reference voltage (Vref) of the new motor by turning the control board trimpot as shown in **(Figure 4b)**. The recommended Vref for this motor is 0.36 volts, although it can vary by model.

**Figure 4.**
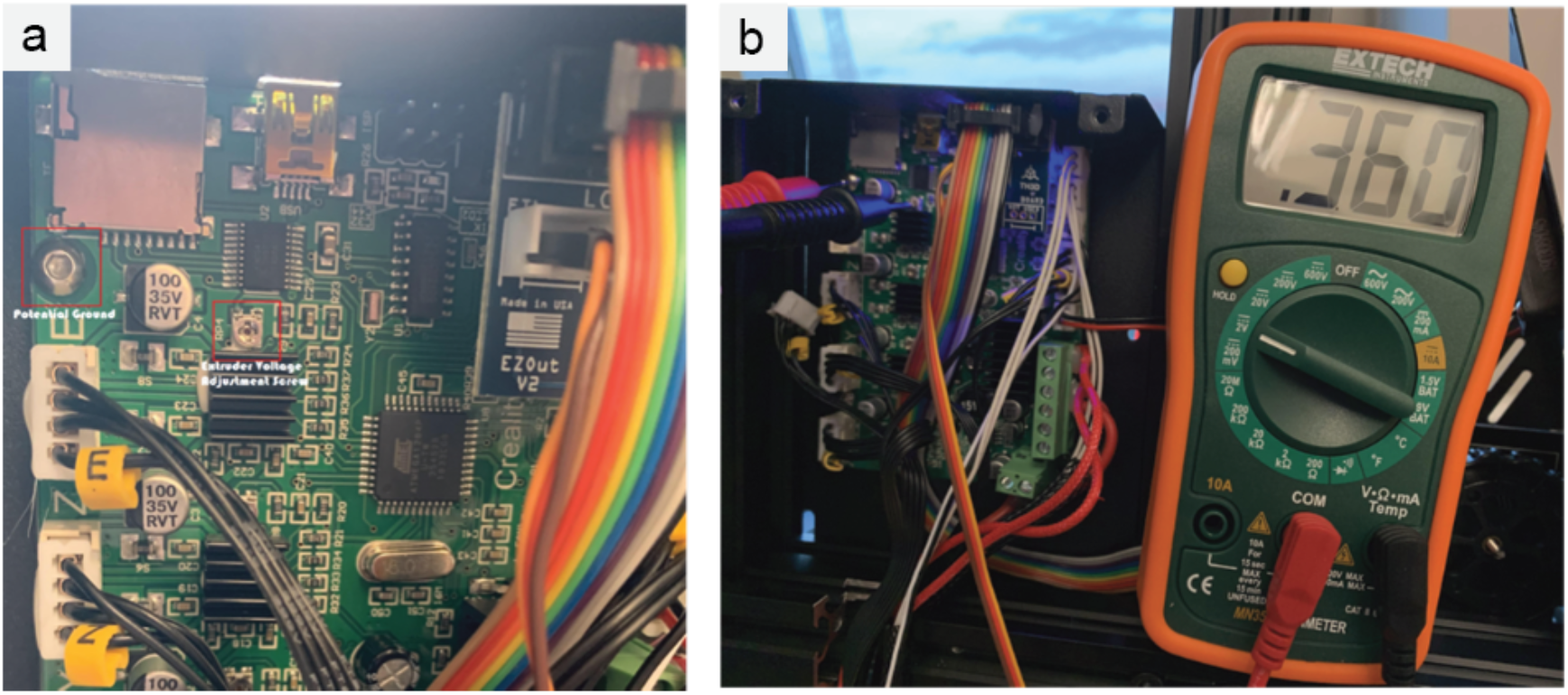
Stepper Motor Voltage Adjustment. (a) The geared stepper motor is plugged into the extruder motor plug on the control board. The reference voltage can be taken using a multimeter to check the voltage from the trimpot when the printer is on. (b) The voltage is adjusted by tightening or loosening the trimpot. For the extruder motor, the Vref should be around 0.36 volts.

### 2.3 Firmware Conversion

3D printing is currently undergoing a gradual shift from 8 bit to 32-bit control boards. 8-bit boards have long been the standard for 3D printers and are capable of performing the tasks needed to run a printer. However, the decreasing cost of 32-bit boards has made them an attractive alternative, and has enabled the development of computationally intense features to take advantage of the increased processing power. At the publication of this paper, 3D printers including the Ender 3 Pro are being shipped with both 8 bit and 32 bit boards. With that in mind, this paper will cover the firmware conversion process for both kinds of board. 32 Bit boards will generally have ARM written on the processor.

The 8-bit firmware used in this project was the TH3D Unified 1 Firmware Package version 1.R2.B5 downloaded from TH3D.com, while the 32 bit firmware was adapted from the TH3D Unified 2 Firmware. These were chosen because TH3D Unified Firmware is an open source firmware project that was designed to be easily implemented, even for those without programming backgrounds. This 8-bit firmware package contains the firmware as well as Arduino IDEs for both Windows and Mac OS X with libraries and detailed setup instructions, and the 32 bit firmware instructions are similarly well documented. This makes these options the most user-friendly method for setting up custom firmware options.

### 2.4 Burning the Bootloader (8 Bit Melzi Board Only)

This step is only necessary when reusing the Melzi 1284p control board that comes with the Ender 3. This is because the firmware shipped on the Ender 3 printer Melzi 1284p control board does not contain a bootloader, preventing firmware changes unless new firmware is burned onto the board. Here, we burn a placeholder firmware that contains a bootloader, allowing firmware changes to be made in the next section. This is done using an Arduino UNO and some DuPont connectors, which costs roughly 6$ **(Table 3)**. This step is not needed if upgrading to an MKS Gen L or using most aftermarket control boards, or with the newer 32-bit control boards. The UNO is connected to the Melzi board through the ICSP headers **(Figure 5a)**, with each header attached to the corresponding pin on both board, with the exception of the reset pin. The reset pin on the Melzi board is instead attached to Pin 10 on the UNO **(Figure 5b)**. The UNO is then connected via USB to the computer to program it to act as a programmer for the Melzi Board. Using the ArduinoIDE downloaded from TH3D, the included Arduino as ISP sketch is loaded into the UNO by opening File->Examples->Arduino as ISP->Arduino as ISP, ensuring that Tools->Board->Arduino Uno and the correct COM port are selected. Then the sketch is uploaded to the Arduino UNO board.

**Figure 5.**
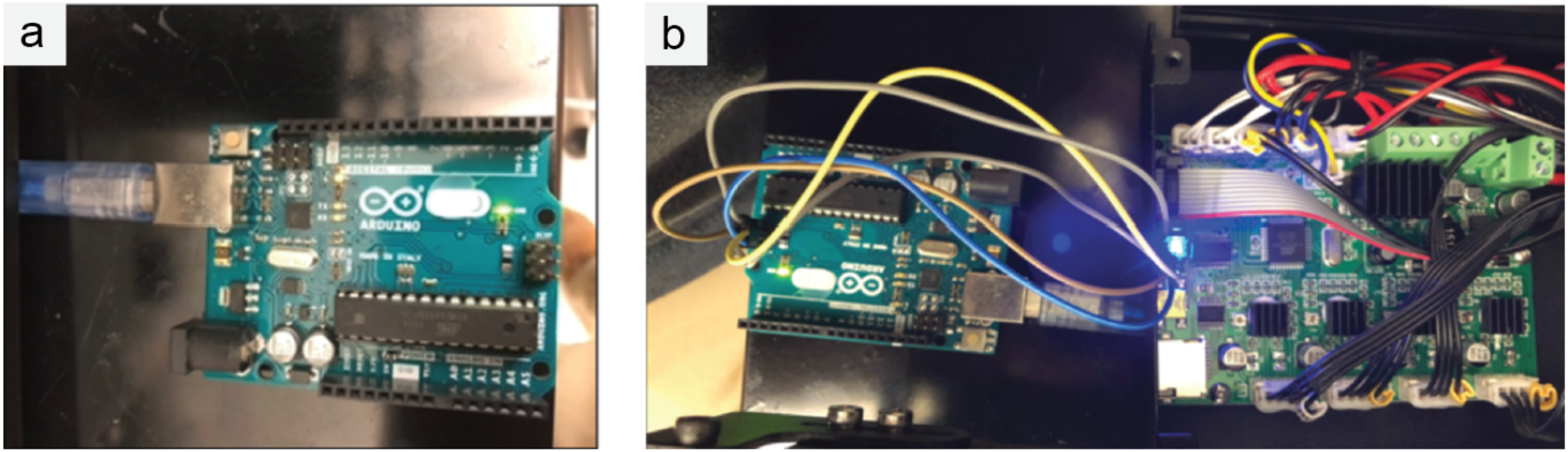
Burning a Bootloader onto a Melzi Board. (a) An Arduino UNO can be programmed with new firmware and used to burn a bootloader onto the Ender 3 control board, allowing further firmware changes to be uploaded. (b) The Arduino ICSP header is connected to the control board ICSP header, except for the reset pin, which is connected to pin 10 on the UNO.

Next, the new bootloader-containing firmware on the Arduino Uno is burned onto the Melzi Board. The Melzi board is selected by choosing Tools->Board->Sanguino 1284p and the Arduino is set as the programmer under Tools->Programmer->Arduino as ISP. This allows the Arduino to act as an in-system programmer that can burn a bootloader onto any AVR board, including the Melzi board in this printer. Then the bootloader is burned over by selecting Tools->Burn Bootloader. Once the bootloader is burned successfully, new firmware can be uploaded directly to the Melzi board via the mini-USB port. This process only needs to be done once for new boards without a bootloader, aftermarket control boards typically come with a bootloader already installed.

### 2.5 8-Bit Bioprinter Firmware Changes

Download & extract the TH3D Unified 1 Firmware package. On computers running OS X, the instructions included in the Mac OS X firmware setup guide were followed. On Windows computers, the OpenFirmwareWindows.bat file can be opened to access the Arduino IDE. Under the configuration.h tab, in the “Creality Ender 3” section of the configuration file, the following changes are made to the file. Note that in Arduino, uncommenting a line is done by removing the two slashes preceding the text.

- uncomment #define ENDER3

The CUSTOM_ESTEPS variable defines how much volume is extruded for each motor microstep. This depends on the stepper motor microsteps per revolution, the thread pitch of the extruder, and the extruder cross-sectional area. Using these factors, we can calibrate the bioprinter to extrude the correct amount of material per microstep.

- Searched & uncommented #define CUSTOM_ESTEPS
- changed CUSTOM_ESTEPS_VALUE to 491

Ordinarily, the firmware prevents extruder motors from operating until the hot end is above a minimum temperature threshold. This command disables this check, enabling printing at low temperatures. Inserted the following line

- #define NO_COLD_PREVENT

The home adjust value is modified to place the new relative position of the nozzle tip at 0,0 when at the home position

- Searched for & uncommented #define HOME_ADJUST
- Defined x and y home adjust locations to (XX) and (YY), respectively

Finally, uncomment #define USER_PRINTER_NAME, and changed the value to a unique identifier to prevent confusion when modifying multiple printers.

If a bed probe is included, the firmware changes for that probe should be performed as well.

### 2.6 32 Bit Bioprinter Firmware Changes

The 32 bit control boards in the Ender 3 Pro, which generally have ARM printed on the processor, require a different firmware, although the customizations made to the actual are very similar. These boards already contain a bootloader and can be overwritten through the USB port. To add custom firmware, the latest version of Python and VSCode are needed, and the Platform I/O Extension should be added within VSCode. Then, the TH3D Unified Firmware 2 Ender 3 Pro Version should be downloaded and extracted. In VSCode, open the folder where the unified firmware was extracted and open configuration.h.

Then:

~~~
Uncomment #define ENDER3_427_BOARD
Type      #define NO_COLD_PREVENT
Uncomment #define CUSTOM_PRINTER_NAME
          #define USER_PRINTER_NAME “CHANGE ME”
Uncomment #define CUSTOM_ESTEPS
          #define CUSTOM_ESTEPS_VALUE 491
~~~

Set the custom Esteps value to 49, so the line should look like

~~~
#define CUSTOM_ESTEPS_VALUE 491
Uncomment #define CUSTOM_PRINTER_NAME
          #define USER_PRINTER_NAME “Example”
~~~

Adding a custom name to the printer will prevent confusion when dealing with multiple bioprinters with different firmwares.

Finally, click the checkmark to compile. The newly compiled firmware will be found under .pio -> build ->

Open this folder in file explorer and copy the latest .bin file to your SD card, insert it into your printer while off, then boot up the printer. Once the printer is booted, reset the firmware to finish the update.

### 2.7 Operating the Bioprinter

At this point, the printer can be operated in basically the same manner as a normal 3D printer. Here, we walk through normal operation. STL files for objects can be created in any CAD program, including open source programs like FreeCAD, OpenSCAD, and Blender. SOLIDWORKS and Fusion360 can also be used, or additional files can be found in online repositories like Thingiverse.

These STL are then imported into a slicer program, Slic3rPE. We use Slic3rPE (Prusa Edition) because it is a free and open source slicer that handles dual extrusion well. This program provides thorough documentation of each setting, making it easy to modify for use with bioinks. Because bioprinting is slightly different than 3D printing, some changes in the slicer settings were needed to slice the STL files for bioprinting. Key settings to change include the extruder and bed temperatures, the infill values, the line width, layer height, and print speed. Bioprinting operates at a lower temperature, so bed and extrusion temperatures were reduced to 37 or set to 0. The nozzle diameters are matched to the diameter of the tapered nozzles, and the layer height set to half of the nozzle diameter. These settings depend on the flow properties of the bioink, with higher viscosity/higher yield stress bioinks able to handle higher layer heights. The infill is set to 100% for most example prints. The complete changes to the Slic3rPE profile are discussed below here.

When Slic3rPE is opened the first time, it activates a setup menu. On this menu, select the Ender 3 Pro as the printer. Under custom printer settings, choose marlin as the printer language. Bed size was set to 235 for x and y. Nozzle size was entered depending on the taper tip gauge in use. this ranged from 22 gauge to 18 gauge, about .4 to .8 mm. Extruder diameter was alternatively set in printer settings for individual nozzles, or under extruder 1 and extruder 2 when using multiple extruders. Filament diameter was left at 1.75 mm. Extrusion temperature and bed temperature were set to 0. Expert mode was chosen to allow all Slic3rPE settings to be available. In the printer settings tab, the number of extruders was assigned depending on the model, either 1 or 2. When using multiple extruders, separate stl files can be added for each extruder to define the print paths of each extruder. Under printer settings, infill was increased to 100% and print solid infill was set to every 1 layer. In print settings, all speeds were changed to 20 mm/s. STL files were then imported and sliced into gcode according to these settings.

Gcode can be sent to the printer using any gcode sender, including pr0nterface, or can be directly read by the printer using the SD card slot on the control board. When using a gcode sender, the correct port and baud rate should be selected for the printer, and slicer programs may need to be closed.

### 2.8 Dual Extrusion Hardware

The hardware conversion is largely similar to a single extrusion conversion, except that the dual extruder syringe bracket assembly is printed instead of the single extruder syringe bracket assembly, and a separate housing for the control board also needs to be printed. 2 motors, 2 rails, 2 motor brackets, 2 motor adapters, 2 barrels, and 2 nozzle adapters are all needed, as well as dual heating pads and thermistors when both barrels were to be thermally regulated. Because the Melzi board that comes with the printer has no second port to connect another extruder, a new control board is necessary to control both extruders.

Here we used an MKS Gen L control board, which can be ordered for about 15$. This board has 5 stepper motor outputs, each of which requires a stepper motor driver module. We used A4988 stepper motor driver modules, which can be purchased for about 1.60$ each at current prices. The voltages on these boards are adjusted via trimpots **(Figure. 4b)** and should be set to 0.8V for X, Y, and E motors, and 0.9V for the Z motor. This board also features additional heater outputs and thermistor inputs needed to regulate temperature in the additional extruder, and superior processing power and memory over the original Melzi board. MKS Gen L boards also already contain a bootloader, so there is no need to flash one onto this board as described above for the Melzi board that came with the printer.

MKS Gen L boards have a larger footprint than the Melzi board, making a different housing necessary for the control board. For this purpose, we printed a new MKS gen L board housing using PETG. This housing replaces the original housing used for the Melzi board, and reuses all the same wires and fans from the original. The wires were carefully labeled as they were transferred to the new board.

### 2.9 8-Bit Dual Extrusion Firmware

Once the board was securely installed and wired, firmware was uploaded to the new control board. MKS Gen L boards do not require a bootloader, so firmware installation proceeded by simply uploading new firmware through the micro-USB port on the board. As noted in the firmware, additional support for this step can be found at http://mksguide.th3dstudio.com.

Dual extrusion requires a few more changes to the firmware than the single extruder setup. Changes are made under the configuration.h tab in the MKS_PRINTER section of the configuration file that will allow you to configure the new board as detailed here. Note that uncommenting each line is done by removing the two slashes that precede the text.

- Uncomment #define MKS_PRINTER

Set the following variables

- MKS_X_SIZE 235
- MKS_Y_SIZE 235
- MKS_Z_SIZE 250
- MKS_X_DRIVER A4988
- MKS_Y_DRIVER A4988
- MKS_E0_STEPS 491
- MKS_E1_STEPS 491
- Uncomment #define DUAL_EXTRUDER_DUAL_NOZZLES
- If keeping the standard display option rather than the 12864 LCD with an SD card, also uncomment define CR10_STOCKDISPLAY

Next, search for DUAL_ HOTEND_X_DISTANCE and then set its value to 30. This defines the distance between the extruder tips, allowing the printer to accurately swap between the two.

- Beneath this line, enter the following:
  #define NO_COLD_PREVENT

Ordinarily, the firmware prevents extruder motors from operating until the hot end is above a minimum temperature threshold. This command disables this check, allowing printing at low temperatures. A custom name for your printer is also recommended. Search for USER_PRINTER_NAME and replace the “CHANGE ME” with the preferred printer name. These changes can also be combined with either bed probe, if desired.

#### 2.9.1 Dual Extrusion 32-bit Firmware

Dual extrusion can also be implemented using a 32-bit board by broadly following the same steps outlined in section 2.6. Briefly, download the TH3D Unified Firmware 2 for MKS Gen L, and open VScode with the PlatformIO extension as outlined in section 2.6. In the configuration.h tab,

-Uncomment #define E1_STEPS_MM
-Set E steps for both motors to 491
-Set the distance between the nozzles to 30 mm by defining
  #define HOTEND_OFFSET_X { 0.0, 30.00 }
-It is also recommended to set a printer name with the lines
  #define CUSTOM_PRINTER_NAME
  #define USER_PRINTER_NAME “Your Name Here”
-In the configuration_backend file, comment out
  PREVENT_COLD_EXTRUSION
-Define EXTRUDE_MINTEMP as 0

This setup was tested using an MKS Gen L board using an MKS Gen L 32 bit board with TMC2209 stepper drivers in UART mode.

### 2.10 Dual Extrusion Use Notes

It is critical to ensure that ensure that both nozzles are perfectly level with each other. Both nozzles can be adjusted slightly by tightening or loosening the luer-locks to ensure that they are precisely level. In SLIC3RPE, dual extrusion prints are designed by loading two separate STL files and assigning an extruder to each one. Because the offset between nozzles is already set in the firmware, it does not need to be set in the slicer program. If adjustments to the offsets between the nozzles are required, they can be adjusted in the slicer or directly in the firmware. The firmware and slicer offsets are additive, so adjustment to both should add up to the appropriate value. The dual extruder setup was utilized to 3D print a customized Texas A&M University logo which was printed using different colored inks in each head (**Figure 7**).

### 2.11 Thermoregulation

This section describes how to regulate barrel temperatures by adding a wraparound heating element to the extruder barrel. This process can be done without altering the firmware by reusing the existing heater ports and thermistor. The hot end heater can be replaced with a heater pad and secured around the barrel with heat shrink tubing. This process uses a 45×100mm 24V flexible polyimide adhesive heater pad, 1” to 0.5” heat-shrink protective tubing, and fully insulated heat-shrink quick disconnect terminals for 22-gauge wire. The extruder barrels are cleaned with alcohol, then the adhesive heater pad is wrapped around the barrel. The thermistor is then laid on top of the heater pad, and the heat shrink tubing shrunk around both to protect the heater and thermistor while holding them in place. The hot end heater wires were cut and spliced into the female 22 gauge quick disconnect terminals, while the heater pad wires are connected to the male quick disconnect terminals. The thermistor wires were cut & reconnected using JST-XH connectors. Care was taken to splice cleanly to ensure that thermistor performance was not affected. These connections allow the barrel to be detached & reattached quickly from the printer without requiring access to the control board. The temperature of the barrel can be set in the slicing program or later in the printer control program by setting the extruder temperature to the desired value.

### 2.12 Bed leveling

Bed leveling probes are used in 3D printing to set a more precise distance between the print bed and nozzle than can be set using traditional endstops. They work by mounting onto the printer carriage and detecting proximity to the print bed, either via mechanical probing or through a capacitive sensing. Bed leveling probes can be also be used to map out the print surface, allowing the printer to compensate for slight variations bed height. These properties make bed sensing probes particularly useful for bioprinting, since bioprinted materials can be highly sensitive to minute variations in bed height. However, due to their position near the extruder, bed probes can interfere with bioprinting in suspension mediums, petri dishes, and well plates. Therefore, it is included here it as an optional addition. Both mechanical and capacitive probes were tested, as each has different advantages. Capacitive probes are completely closed and have no moving parts, which makes them more practical to sterilize than mechanical probes. However, capacitance can vary with surface materials and temperature, so the probe must be recalibrated if the print surface or ambient temperature changes significantly. On the other hand, mechanical probes have moving parts and exposed circuitry, but detect surfaces regardless of material or ambient conditions.

Bed leveling kits were purchased separately, the two we tested were the ANTCLABS BLTouch(mechanical) and the TH3D EZABL Pro(capacitive). The BLTouch is a mechanical probe that costs about 40$, while the EZABL is a capacitive probe costing roughly 65$. Both were purchased directly from suppliers to avoid common counterfeits. For both the BLTouch and the EZABL, separate syringe bracket assemblies were created with mounting points for the probes, which are included in the supplementary files.

### 2.13 BLTouch Hardware

The BLTouch is installed onto the mounting point on the SBA using M3 screws **(Figure 6a)**. The BLTouch was installed using a pin 27 board, which separates 5V, ground, and signal pins from the LCD header to be used for the BLTouch **(Figure 6a)**. These 3 pins on the BLTouch extension cable can be connected here, care should be taken that the wires are correctly ordered. The 2-pin connector replaces the Z-endstop on the control board. Both the 2-pin and 3-pin connectors can be secured in place with hot glue or replaced with JST connectors to prevent loosening over time. The extension cable is then pushed through the wire sleeve to the endstop and connected to the BLTouch.

**Figure 6.**
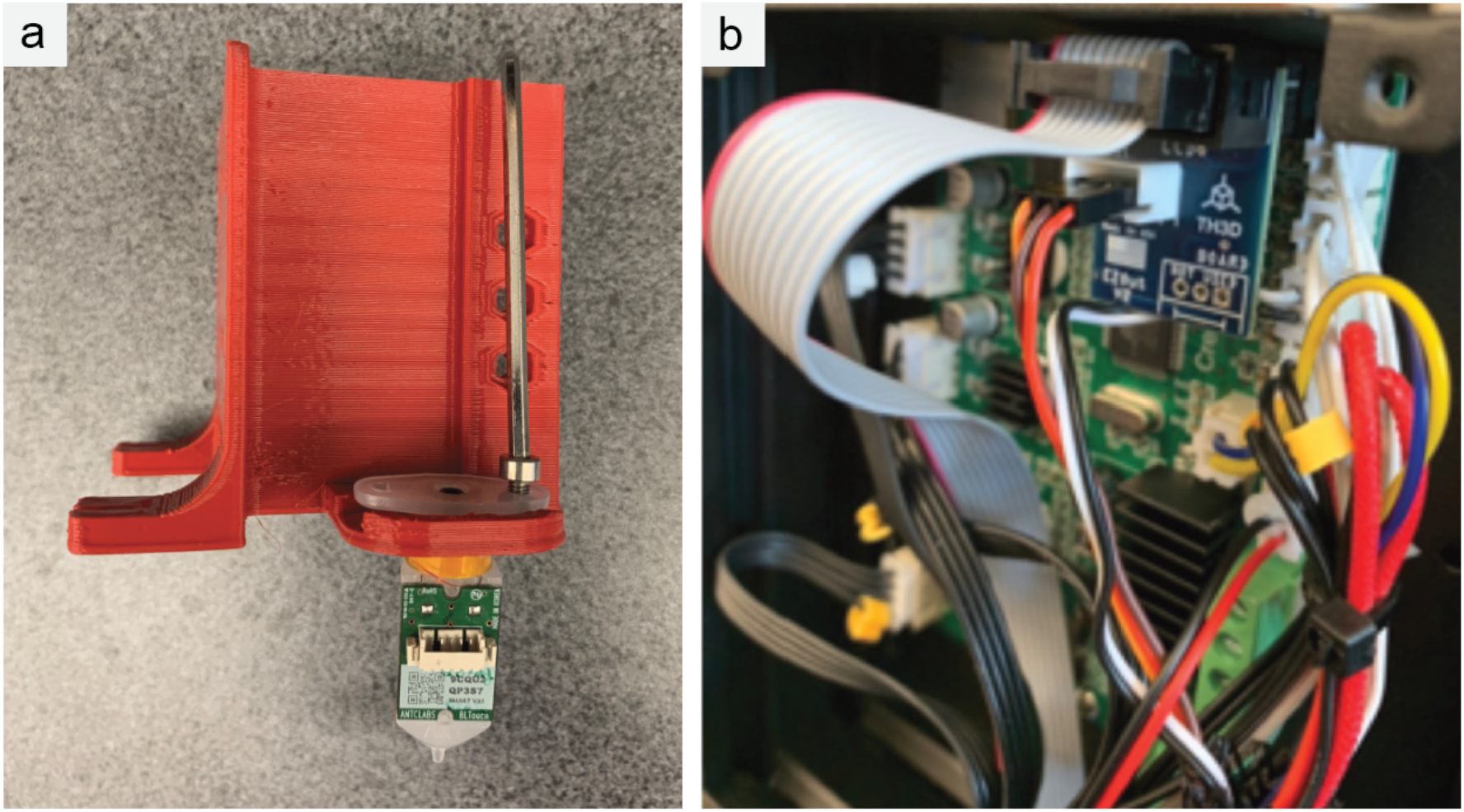
BLTouch Installation. (a) The BLTouch is mounted into the printed attachment point on the syringe bracket assembly using M3 bolts and nuts. (b) The Pin27 board provides a separate pinout from the LCD header to be used for BLTouch installation.

**Figure 7.**
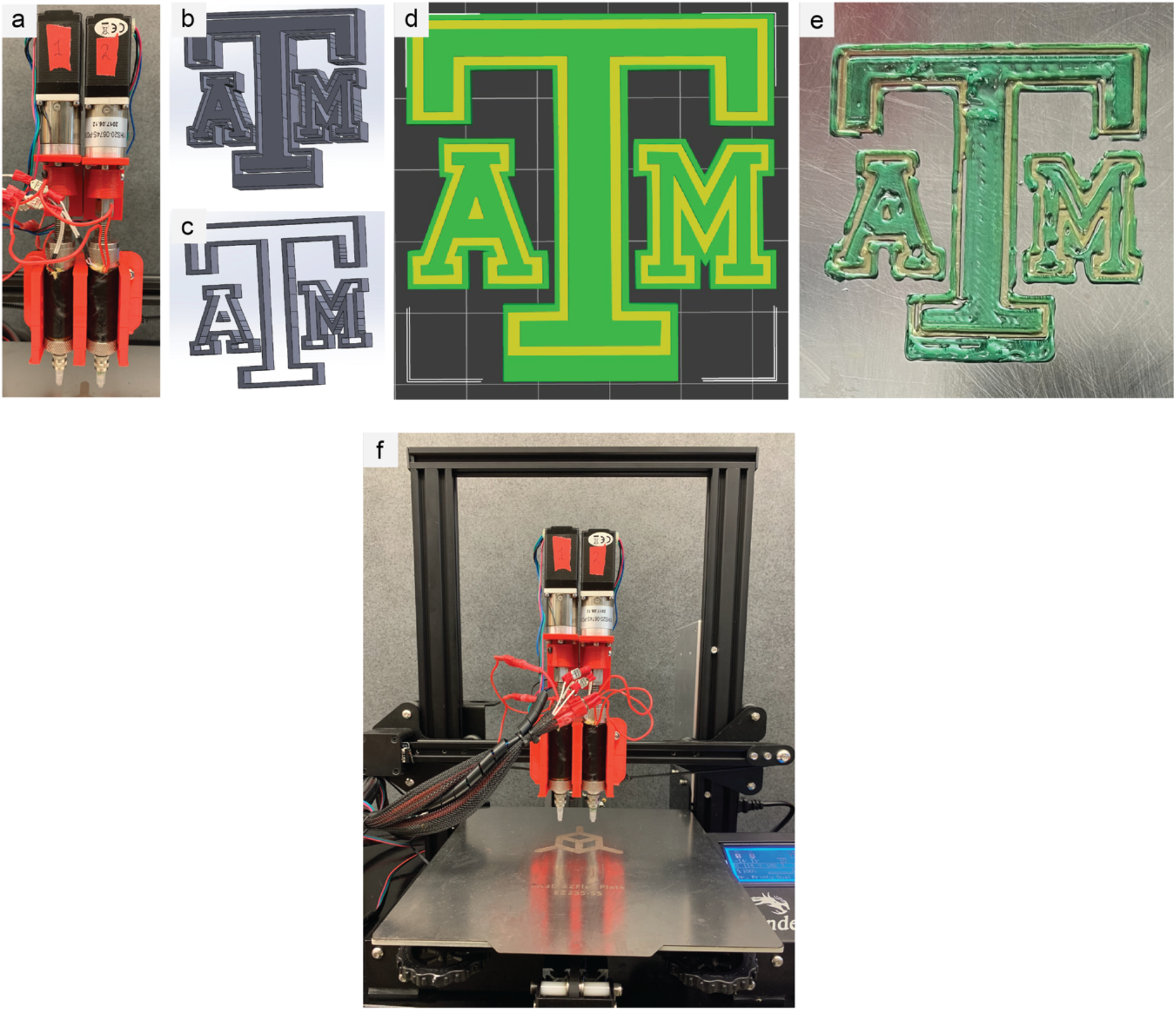
Dual Extruder modification and Multimaterial 3D Printing. (a) Dual extruder bracket with extruders attached. (b) .STL file that extruder one will print. (c) .STL file that extruder two will print. (d) Both .STL files imported to PrusaSlicer. (e) Final dual extrusion 3D printing. (f) The customized dual extruder 3D bioprinter.

**Figure 8.**
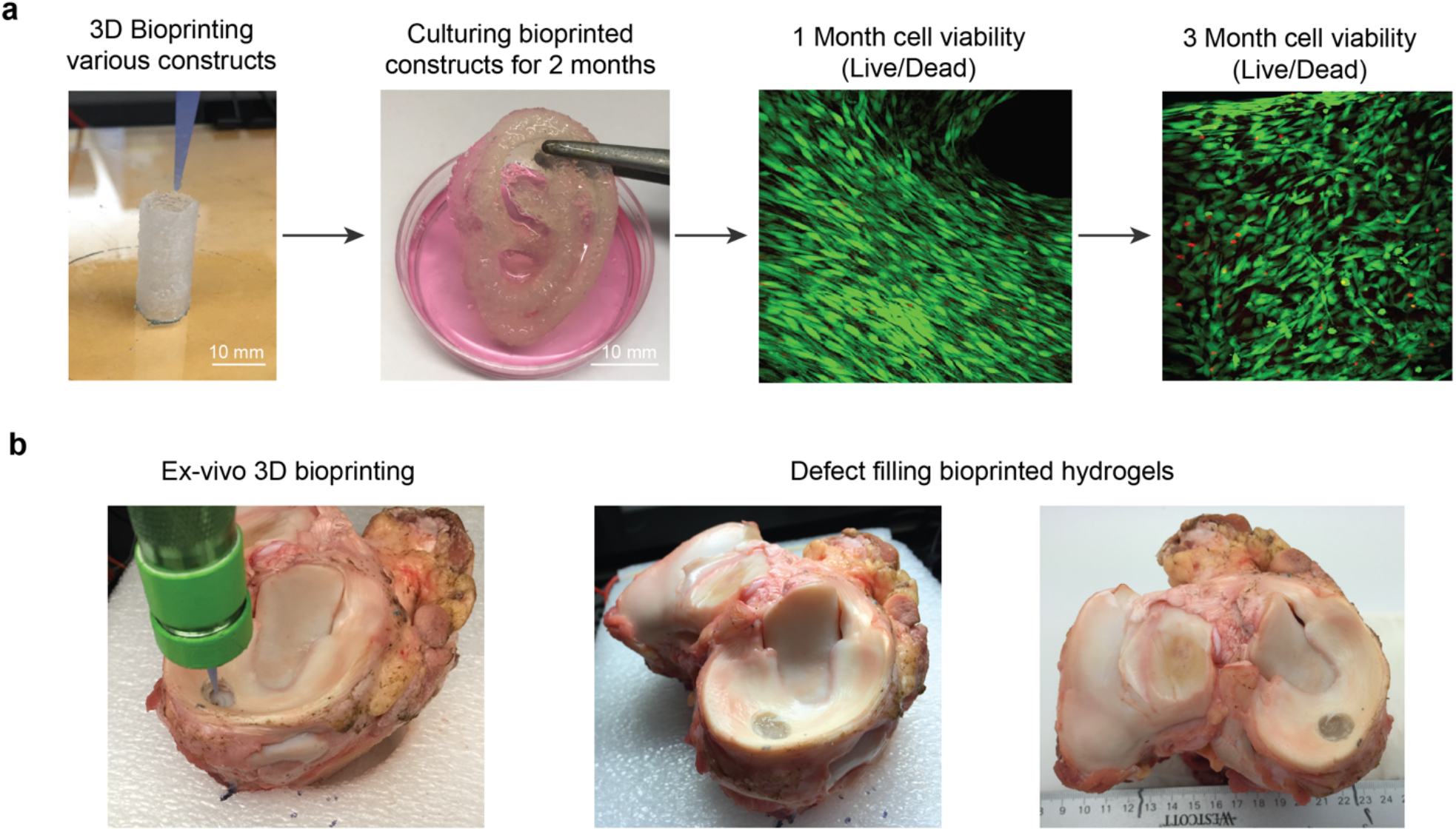
Demonstration of Practical Utility for 3D Bioprinting. (a) Bioprinted constructs exhibit 3D printability with NICE bioinks. Encapsulated 3T3 murine preosteoblasts proliferated *in vitro* and maintained high viability for over 90 days. (b) *Ex vivo* bioprinting into an induced meniscal defect demonstrated excellent adhesion to surrounding tissue even through repetitive stress cycles. These results suggest that in vivo bioprinting of NICE bioinks into damaged cartilage regions may have potential for future tissue engineering treatments.

### 2.14 BLTouch Firmware

These firmware changes were made in addition to the changes in the firmware for either the single extruder or dual extruder. In the configuration.h tab, search for BLTOUCH and uncomment the lines #define BLTOUCH and #define SERVO0_PIN 27. Then, uncomment the line #define CUSTOM_PROBE in either the Ender 3 Section for single extruder setups or the MKS GEN L Section for multi-extruder setups. Then search for X_PROBE_OFFSET_FROM_EXTRUDER and set to the measured offset between the probe and the primary nozzle in the x direction, and Y_PROBE_OFFSET_FROM_EXTRUDER should be set to the y offset. Finally, the printer was tested to ensure functionality, and the Z-probe offset was measured and set in the printer menu.

### 2.15 EZABL Hardware

As with the BLTouch, the EZABL was installed on the appropriate SBA mount, and was installed according to the detailed manufacturer instructions. Briefly, the EZABL height relative to the extruder was set to 2mm above the extruder tip using the included retaining nuts and blue Loctite. The EZABL Control board was mounted on the printer body near the power supply, and the power wires were directly connected to the power supply, while the z-endstop wires were connected from the 3D printer control board to the EZABL control board. Next, the EZABL sensor plug was connected into the EZABL control board, and the install was checked by testing the capacitive sensor by fingertip.

### 2.16 EZABL Firmware

These firmware changes were made in addition to the changes in the firmware for either the single extruder or dual extruder.

Uncomment the line #define CUSTOM_PROBE in either the Ender 3 Section for single extruder setups or the MKS GEN L Section for multi-extruder setups. Then search for X_PROBE_OFFSET_FROM_EXTRUDER and set to the measured offset between the probe and the primary nozzle in the x direction, and Y_PROBE_OFFSET_FROM_EXTRUDER was set to the y direction offset. The Z-offset was then set according to the manufacturer instructions by adjusting the calibration screw for the appropriate sensitivity and setting the z-offset via the printer menu interface.

### 2.17 3D Bioprinting

Bioprinting was carried out using a Nanoengineered Ionic Covalent Entanglement(NICE) bioink composed of 10% w/v gelatin methacryloyl, 1% w/v kappa carrageenan, 2% Laponite XLG nanosilicates, and 0.25% Irgacure 2959 in DI water.^16^ MC3T3 Subclone 4 (ATCC CRL-2593) cells were incorporated into the bioink at a density of 3.33 × 10^6^ cells/mL by resuspending a centrifuged cell pellet in 200 μL of Dulbecco’s Modified Eagle Medium with 10% Fetal Bovine Serum, and gently mixing with the bioink. The bioink was kept at 37 °C prior to extrusion.

Bioprinting was carried out through a 400 μm tapered Luer lock tip, line width was set to 500 μm, layer height to 200 μm, and print speed at 0.15 mL/minute (20 mm/s). The printed hollow cylinders were 2 cm tall with an outer diameter of 10 mm and an inner diameter of 8 mm. Live/dead staining was used to carry out cell viability tests by incubating scaffolds in 1 μL/mL calcein AM and 2 μL/mL ethidium homodimer for an hour, followed by soaking in PBS to remove non-absorbed strain. Images were taken using a confocal microscope to evaluate cell viability both at the surface and at depth **(Figure 8a)**.

*Ex vivo* bioprinting tests were performed on an equine upper tibia section with attached meniscus cartilage, donated from the Texas A&M Large Animal Hospital. A power drill was used to induce a 1 cm defect in the meniscal cartilage. The tibia was secured to the print bed using a styrofoam placeholder. The defect size and position were modeled in Prusa Slic3r and Pr0nterface. After printing, the bioink was crosslinked using 0.25 mW/cm^2^ of UVA light for 60 seconds, and a potassium chloride solution was sprayed over the bioprinted area. The bioprinted implant was subjected to cyclic mechanical compression to ensure good adherence to the native meniscal cartilage **(Figure 8b)**.

## 3. CONCLUSION

Despite the burgeoning popularity of bioprinting, its adoption has been slowed by the high cost of bioprinters. Restrictive user interfaces and proprietary hardware also drive-up costs associated with bioprinting, in contrast to the largely modular and open-source field of 3D printing. In this paper, we demonstrate a method for converting a thermoplastic printer into a bioprinter. This conversion process allows bioprinting to incorporate the modularity, lower costs, and open-source software available for mainstream 3D printers that have developed over the last decade thanks to the enormous popularity of additive manufacturing.

The bioprinter conversion process we detailed in this paper contains several key innovations. Firstly, the provided 3D models and firmware make this conversion possible in only a few hours for about 400$. This low-price tag is comparable to many hobbyist 3D printers today, making bioprinting as accessible as 3D printing. Further, most of the 3D printer does not need to be modified for this conversion, and the 3D printer itself is used to print most of the necessary hardware. By basing this printer on popular 3D printing hardware, this bioprinter can be customized with a wide array of modular add-ons. As a bioprinter, the printer can continue to run through software designed for thermoplastic printers, allowing use of open source software that enables the bioprinter to be customized for any research needs. To demonstrate the customizability of this setup, we have included options for single and multi-extruders, temperature control, and two bed leveling options. By including all CAD and firmware files necessary for this conversion, we make this process as streamlined and painless for researchers as possible.

This is also the first publication of a dual extrusion bioprinter run on motor-driven extrusion. This extrusion modality is advantageous for printing advanced bioinks with more complex rheological properties and allows printing a wider range of materials than can be printed using pneumatic extrusion. Most importantly, this paper drastically reduces the cost of the most expensive barrier to bioprinting research, the bioprinter itself. We expect that this publication will help expand the field of bioprinting and contribute to the ultimate goal of biofabrication: creation of functional tissues for clinical use. It is our hope that this paper will contribute to providing new research pathways to cure diseases that currently affect millions of patients worldwide.

## Supporting information

3D Printer Parts

## Acknowledgements

A.K.G. acknowledges financial support from the National Institute of Biomedical Imaging and Bioengineering (NIBIB) of the National Institutes of Health (NIH), Director’s New Innovator Award (DP2 EB026265), National Science Foundation (NSF) Award (CBET 1705852) and President’s Excellence Fund (X-Grants) from Texas A&M University. The content is solely the responsibility of the authors and does not necessarily represent the official views of the funding agency. Some of the images in the article were created with Biorender.

