## Supporting Information for "Designing Cost-Effective, Open-Source, Multi-Head Bioprinters via Conversion of Hobby-Grade 3D Printers"

D. Chimene, K.A. Deo, J. Thomas, Prof. A.K. Gaharwar
Biomedical Engineering
College of Engineering
Texas A&M University
College Station, Texas 77843, USA

Prof. A.K. Gaharwar
Material Science and Engineering
College of Engineering
Texas A&M University
College Station, Texas 77843, USA

Prof. A.K. Gaharwar
Interdisciplinary Graduate Program in Genetics,
Texas A&M University,
College Station, Texas 77843, USA.

Prof. A.K. Gaharwar
Center for Remote Health Technologies and Systems
Texas A&M University
College Station, Texas 77843, USA

**Keywords:** 3D Bioprinting, Open Source 3D Bioprinters, Multimaterial 3D Printing;

**Table S1. Main printer supplies with their purchase information**

| Item | Price | Link |
|---|---|---|
| *Ender 3 Pro* | $209.00 | https://www.creality3dofficial.com/products/creality-ender-3-pro-3d-printer |
| *19:1 Planetary Gearbox NEMA 11 Stepper Motor Extruder 3D Printer Motor* | $40.49 | https://www.amazon.com/19-Planetary- Gearbox-Stepper-Extruder/dp/B00PNEQKOS |
| *ReliaBot 200mm MGN12 Linear Rail Guide with MGN12H Carriage Block for 3D Printer and CNC Machine* | $26.29 | https://www.amazon.com/ReliaBot-Linear- Carriage-Printer-Machine/dp/B07B4F2G48 |
| *Makin's USA Professional Ultimate Clay Gun Extruder* | $36.52 | https://www.amazon.com/Makins-USA- Professional-Ultimate-Extruder/dp/ B004VY6PJG |

**Table S2. Optional Parts**

| Item | Price | Link |
|---|---|---|
| *EZABL$_{TM}$ Pro Mini– Auto Bed Leveling Kit w/ OEM Ender 3 Mount* | $58.99 | https://www.th3dstudio.com/product/ezabl- pro-plug-print-auto-bed-leveling-kit/ |
| *ANTCLABS BLTouch : (With 1M Extension Cable Set)* | $39.29 | https://www.amazon.com/BLTouch-Leveling-Sensor-Premium-Printer/dp/B076PQG1FF/ |
| *Pin 27 Adapter Board* | $10.99 | https://www.th3dstudio.com/product/ezout-v2-filament-sensor-kit-or-bl-touch-adapter-board/ |
| *Steel Build Plate 235mm x 235mm* | $14.99 | https://www.th3dstudio.com/product/ezflex- plate-build-sheet-steel-flex-base/ |
| *USB to Mini-USB data cable* | $4.79 | https://www.amazon.com/UGREEN-Charging-Controller-Players-Receiver/dp/B00P0GI68M |
| *Uno Bootloader Flashing Kit(Arduino Uno, USB 2.0 Cable, M-F & F-F Dupont Wires)* | $15.99 | https://www.th3dstudio.com/product/arduino-uno-bootloader-flashing-kit/ |

**Table S3. Recommended Tools**

| Item | Price | Link |
|---|---|---|
| *Calipers* | $19.99 | https://www.amazon.com/Neiko-01407A-Electronic-Digital-Stainless/dp/ B000GSLKIW |
| *M3 Hex Head Nut & Bolts assortment pack* | $16.99 | https://www.amazon.com/Stainless-Phillips-Machine-Washers-Assortment/dp/B07FCN64HV |
| *Wire Crimpers* | $13.99 | https://www.amazon.com/VISE-GRIP-Stripping-Cutter-8-Inch-2078309/dp/ B000JNNWQ2 |
| *JST-XH Connector Kit with 2.54mm Female Pin Header and 2/3/4/5/6 Pin Housing* | $9.29 | https://www.amazon.com |
| *Size 111 Viton O-Ring* | $8.59 | https://www.amazon.com/dp/B0051Y1VYY |
| *Size 007 Viton O-Ring* | $5.53 | https://www.amazon.com/dp/B000MN9SV2 |
